# RAVEN: development of a novel volumetric extrusion-based system for small-scale Additive Manufacturing

**DOI:** 10.1101/2023.03.30.534759

**Authors:** Pierpaolo Fucile, Vivek Cherian David, Maria Kalogeropoulou, Antonio Gloria, Lorenzo Moroni

## Abstract

Recent technological advances in the field of Additive Manufacturing (AM) and the increasing need in Regenerative Medicine (RM) for devices that better and better mimic native tissues architecture are showing limitations in the current scaffolds fabrication techniques. A switch from the typical layer-by-layer approach is needed to achieve precise control on fibers orientation and pores dimension and morphology. In this work a new AM apparatus, the RAVEN (Robot-Assisted Volumetric ExtrusioN) system, is presented. RAVEN is based on a 7-DOF robotic arm and an FDM extruder and allows for volumetric extrusion of polymeric filaments. The development process, namely the robotic motion optimization, the optimization towards small-scale trajectories, the custom-made hardware/software interfaces, and the different printing capabilities are hereby presented. The successful results are promising towards future advanced applications such as *in vivo* bioprinting, in which the ability of the robot to change its configuration while printing will be crucial.

## 1. Introduction

Regenerative Medicine (RM) is a continuously evolving field, which aims to the regeneration of damaged tissues through the use of living cells, biomaterials, and other therapies [1-3]. One of the key elements of RM is a scaffold, a 3-Dimensional porous structure that catalyzes and enhances the process of regeneration through a delicate cell-material interaction [1, 4]. Indeed, tissue regeneration occurs since scaffolds main requirement is to mimic the Extra Cellular Matrix (ECM) of the native tissue, in order to send a series of specific biological signals that guide the process of cell adhesion, proliferation, and differentiation, so to create new tissue in place of the damaged one [4, 5]. To achieve this, there are many chemical and structural requirements that scaffolds must satisfy, such as (i) biocompatibility, (ii) biodegradability, and in particular a tailorable degradation process comparable to the typical regeneration times of the native tissue, (iii) a fully interconnected pores network, which allows for (iv) adequate mass transport properties during the regeneration process, and (v) load bearing function [6]. The need for a tailored device in terms of structural features led the field towards more complex fabrication methodologies, namely Additive Manufacturing (AM) techniques, also known as “3D Printing”, which allow for the production of devices with a high degree of customization in a layer-by-layer fashion [5].

In the field of RM, these techniques allow for the fabrication of customized scaffolds with a high degree of interconnection in the inner porous structure [3, 5], and replaced more “conventional” techniques (e.g., solvent casting, particulate leaching, freeze-drying, gas foaming), which provide a limited degree of interconnection of the pores, lacking control over pores dimension and geometry as well as over long-range microarchitecture.

However, as researchers aim to an always increasing degree of complexity (i.e., design-wise) towards a better mimicking of native tissues, limitations in AM, especially for extrusion based systems, are evident when it comes to the fabrication of devices with complex architectures (i.e., anisotropic structures and specific fibers disposition).

In standard AM, the printing extruder moves in a Cartesian-linear fashion, layer after layer. Thus, as the thickness of each layer is fixed, it would seem more correct to talk about “2.5D Printing”, which is then repeated to create a volumetric 3D object [7]. Furthermore, the human body is not built layer-by-layer. Cells and proteins are synthesized and organized in a volumetric manner.

In this context, a shift from the typical x-y layer-by-layer fabrication approaches to a more “volumetric” AM is crucial. In order to fabricate 3D scaffolds for RM with enough structural complexity, a novel AM technology with extra Degrees of Freedom (DOF) is needed. New systems with more DOFs have already been implemented in large-scale manufacturing (e.g., industrial and manufacturing fields) [8-10] and more complex structures were fabricated in a “less planar” fashion. In particular, robotic arms have been implemented in the process, allowing for higher control (i.e., orientation of the printing head in a desired way to print volumetrically). Such an approach is quite common in the construction field, in which concrete and clay-like materials are extruded with the support of robots [11-14].

Few attempts towards robotic AM implementation have been made in small-scale manufacturing. Some applications involved multi-axis motion, in which the combination of the robot’s DOF and a tilting printing bed allowed for nonplanar layers deposition [15-17]. Furthermore, some advances were also made towards non-planar printing, which has been achieved through the extrusion of non-planar surfaces obtained as result of custom-made non-planar slicing processes [17-19], or through BREP-based helical/tubular structures [20]. These approaches are very promising, though far from an actual polymeric volumetric printing. However, when it comes to RM, and thus small-scale manufacturing with medical grade polymers, this application is more challenging. Some first attempts have been made in that direction [21, 22], though with soft/curable materials and simple architectures. The introduction of additional DOF in scaffolds fabrication can significantly extend the capabilities of AM in terms of change of extrusion direction during the process as well as the deposition of complex nonplanar layers. In the field of biomedical applications this would result in many advantages, such as the possibility of *in situ* AM as well as of the fabrication of supportless structures and more mechanically reliable scaffolds (i.e., mechanical properties are affected by the precise alignment of fibers).

In this context, the aim of this paper is to present a new custom-made universal extrusion-based AM system named RAVEN (Robot-Assisted Volumetric ExtrusioN), which is based on a 7-DOF robotic arm and will be crucial in the development of advanced small-scale fabrication of scaffolds for RM. The innovations proposed in this approach are many. First of all, it is based on a 7-DOF robotic arm. Compared to a 5 or 6-DOF robotic arm which is widely used in the robotic/manufacturing field [12, 23], the 7^th^ or *redundant* DOF improves the outcome of the whole process. The robot can change its configuration while keeping the end-effector on the same position, and this paves the way to advanced obstacle avoidance approaches during printing and complex layers deposition. Though, this increases the complexity of the whole system as well [24, 25]. From a mathematical point of view the solution of the Inverse Kinematic (IK) problem, which is the basis of every robotic approach, is much more difficult. In general, the end-effector (and thus, the extruder) velocity is related to the other joints’ rotational velocities through the determinant of the Jacobian matrix. In order to control the robot over specific trajectories, the joints’ rotational velocities must be calculated from the inverse Jacobian matrix [26, 27]. Such operation is very difficult when the matrix is not symmetric, as in the case of 7 DOF. In this context, *ad hoc* control algorithms must be developed to properly control and exploit the potential of the redundant DOF.

Indeed, the RAVEN AM system implements custom-made control algorithms, both for the trajectory planning/execution and the communication with the extruder, which occur simultaneously. This is why this system aims to be a *universal platform*, which can be easily adapted to different robots (also with less DOF) and extruders. This new system allows for the extrusion of continuous lines in air, thanks to the implementation of an advanced cooling system that allow for a fast cooling of filaments without support structures. Compared to other systems, in which only segments can be printed, this prevents defects in the final results (e.g., in case two segments do not attach properly), and points of discontinuity. First trials towards RM showed the capability of the RAVEN system in terms of change of printing directions/angles, cooling effectiveness and volumetric printability without over-extrusion.

## 2. Materials and Methods

### 2.1 Hardware Setup

The RAVEN system hardware is shown in Figure 1. The robotic arm is a 7-DOF Sawyer Robotic Arm (Rethink Robotics GmbH, Germany). A traditional Fused Deposition Modelling (FDM) extruder (V6 Hemera XS, E3D, Oxford, Oxfordshire) with a 0.4 mm needle diameter is employed and attached to the end effector of the robot through custom-made and tailored supports, which were designed in SolidWorks 2022 (Dassault Systemés, France) and then 3D Printed through FDM. Thus, the concept of RepRap (Replicating Rapid Prototyper), in which a 3D printer is made with 3D printed parts, is central [28].

**Figure 1:**
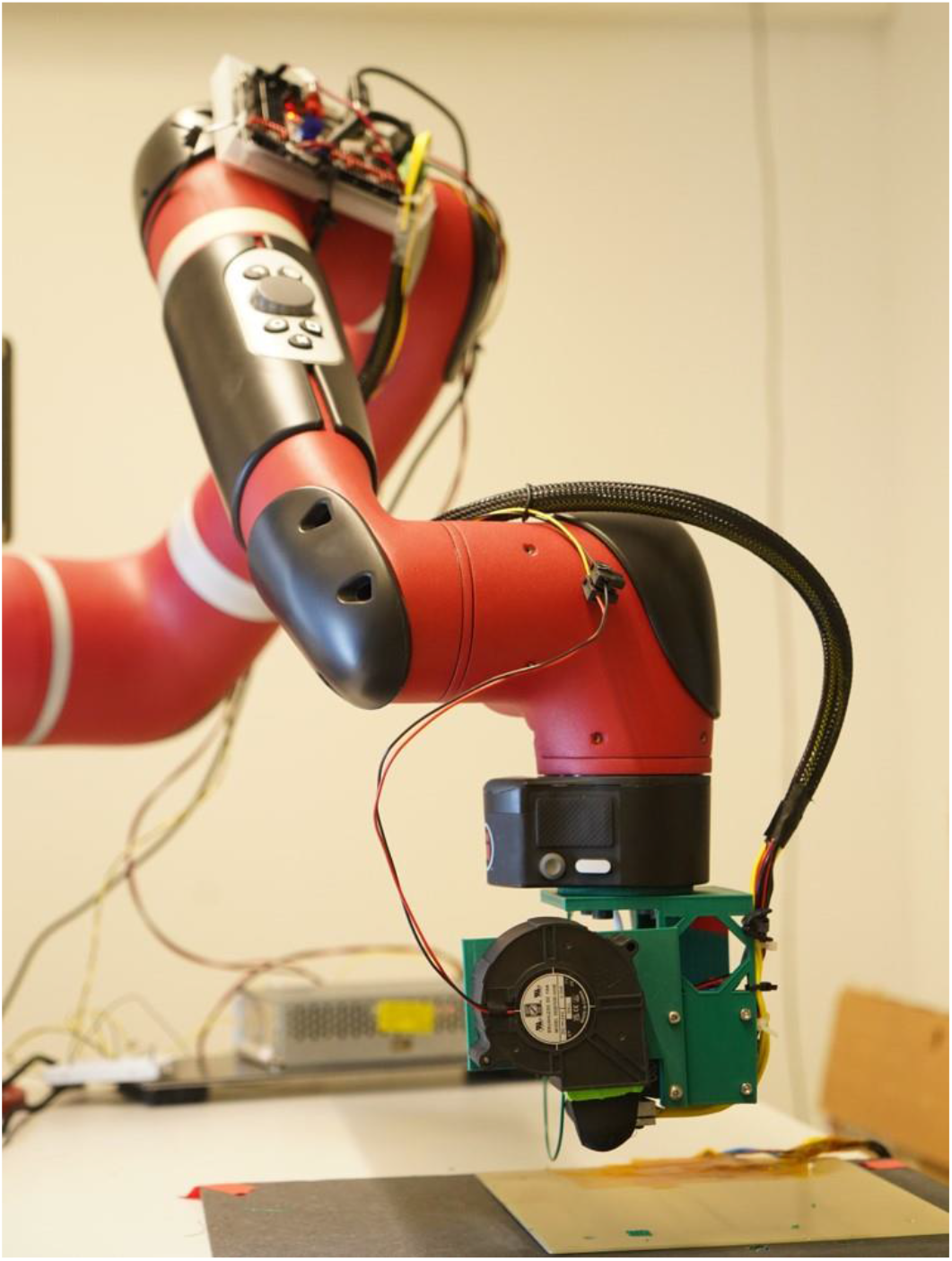
The RAVEN system hardware.

The extruder is controlled through a 32-bit controller board (BigTreeTech SKR 2, BigTreeTech, China). The cooling flow is provided by a traditional fan for 3D Printing (RS PRO Centrifugal Fan, RS Components B.V., Netherlands), which is attached to the extruding system through custom-made and tailored supports, which were designed in SolidWorks 2022 as well. Since specific directions of the air flow are needed, *ad hoc* nozzle fan duct were designed and printed.

### 2.2 Optimization of motion planning

In order to achieve a precise control on robot movements towards volumetric fabrication, motion optimization was performed [25]. The robot was controlled within the Robotic Operating System (ROS), which is an open-source linux-based environment that allows for communication through a series of built-in and customizable packages and libraries, and MoveIt (PicNik Robotics, Colorado, USA) that is an open-source platform for robotic motion planning, manipulation, perception, and control. For a given list of waypoints (i.e., the desired trajectory) that the robot has to follow, the movement is a combination of planning and IK solving. Thus, different approaches were tested in terms of planners and solvers. For small-scale AM purposes, Cartesian trajectories deriving from cartesian points must be generated and precisely followed by the robot. In this context, different combinations of the two were implemented. In particular, the Kinematics and Dynamics Library (KDL) solver (Orocos Project) was coupled with the MoveIt Cartesian Planner and PILZ Industrial Planner, both provided by the MoveIt libraries, while the IKFast IK Solver (OpenRave, Robotics Institute, Carnegie Mellon University, Pittsburgh, PA) was selected to be paired to the Descartes Planner (ROS-Industrial). MoveIt Cartesian Planner – KDL and PILZ Industrial Planner – KDL were selected as they are quite common in the robotics community, when it comes to 5 or 6-DOF robots in the industrial and automation field. While Descartes Planner and IKFast IK Solver were selected by the authors as they are some of the best choices for small motions along Cartesian trajectories and towards 7 DOF robotic movements.

The different approaches were tested on the same desired cylindrical trajectory, as shown in Figure 2. The outcome of each trial was a plot of the desired trajectory vs. the robot’s end-effector joint actual movements, which were recorded exploiting ROS capabilities through a custom-made code. Attention was given to the capability of the different techniques to have full control on crucial motion parameters (i.e., velocity and acceleration). Every trial and coupling were first simulated in Gazebo (Open Robotics, Mountain View, CA, USA), which is a simulated robotic environment. Towards the final application of RAVEN system, the Descartes Planner and IKFast IK solver were selected.

**Figure 2:**
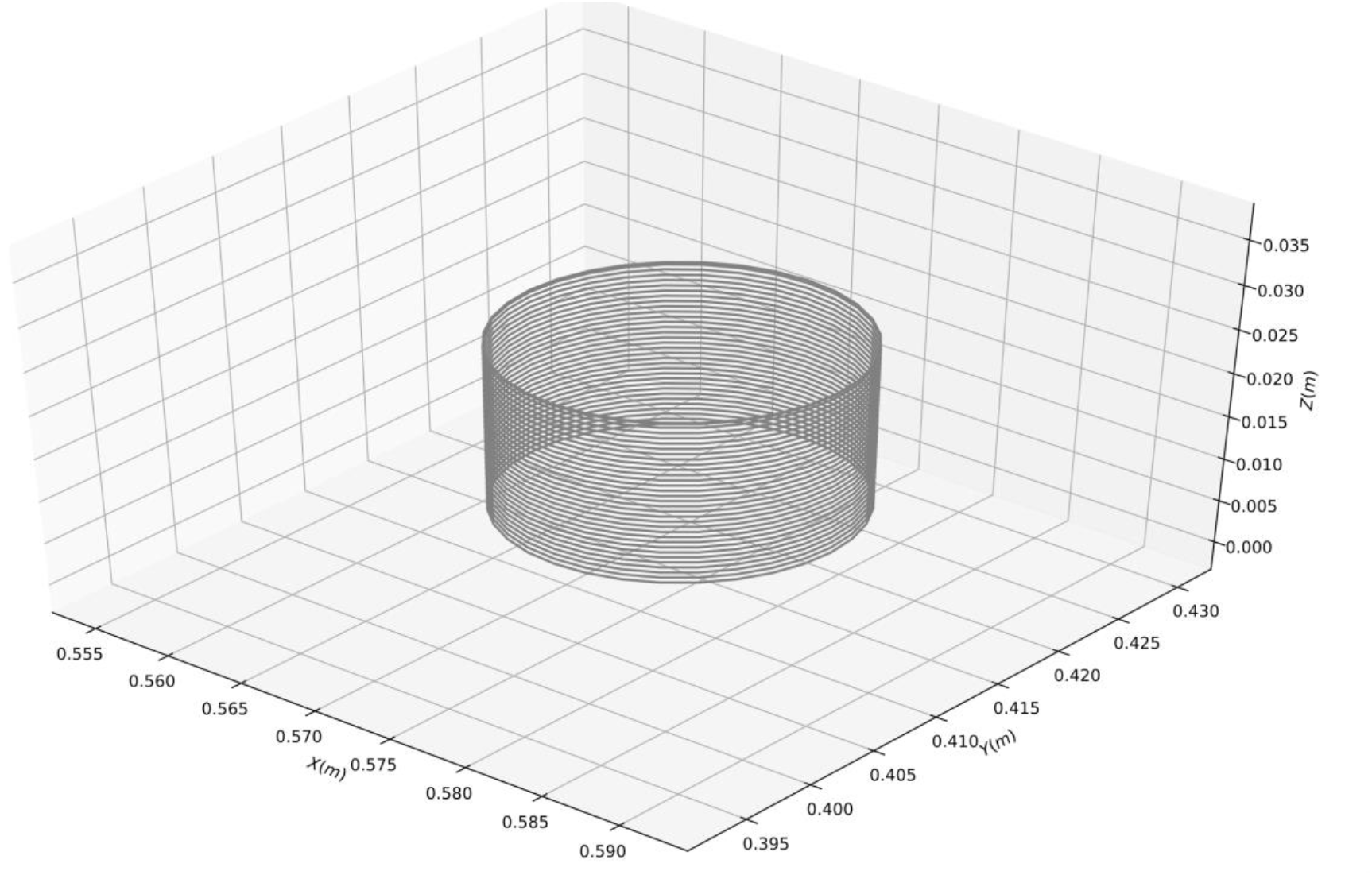
The cylindrical trajectory generated for motion planning optimization.

### 2.3 Trajectory generation and planning

The selection of the best planner and solver towards the specific application, namely Descartes Planner and IKFast IK solver, led to the development of a custom-made platform for the precise control over the Sawyer robotic arm motion towards small-scale motions for AM. Many tests were conducted to optimize the code and the accuracy of the trajectory generated by the new planner, Descartes. The path planning system was optimized in order to improve its efficiency. A complex trajectory made of horizontal and vertical segments (Figure 3) was generated point by point and tested at different motion velocities, namely 0.4, 0.8, 0.32, 1.6, 2.4, 3.2, 4.0, 8.0, 16.0, 21.0, 32.0, 40.0, 80.0, and 160.0 mm/s. Furthermore, each segment was characterized by different points density in order to validate the behavior of the trajectory generation algorithms in different conditions, namely 34, 3, and 5 points for segments #1, #2, and #3, respectively.

**Figure 3:**
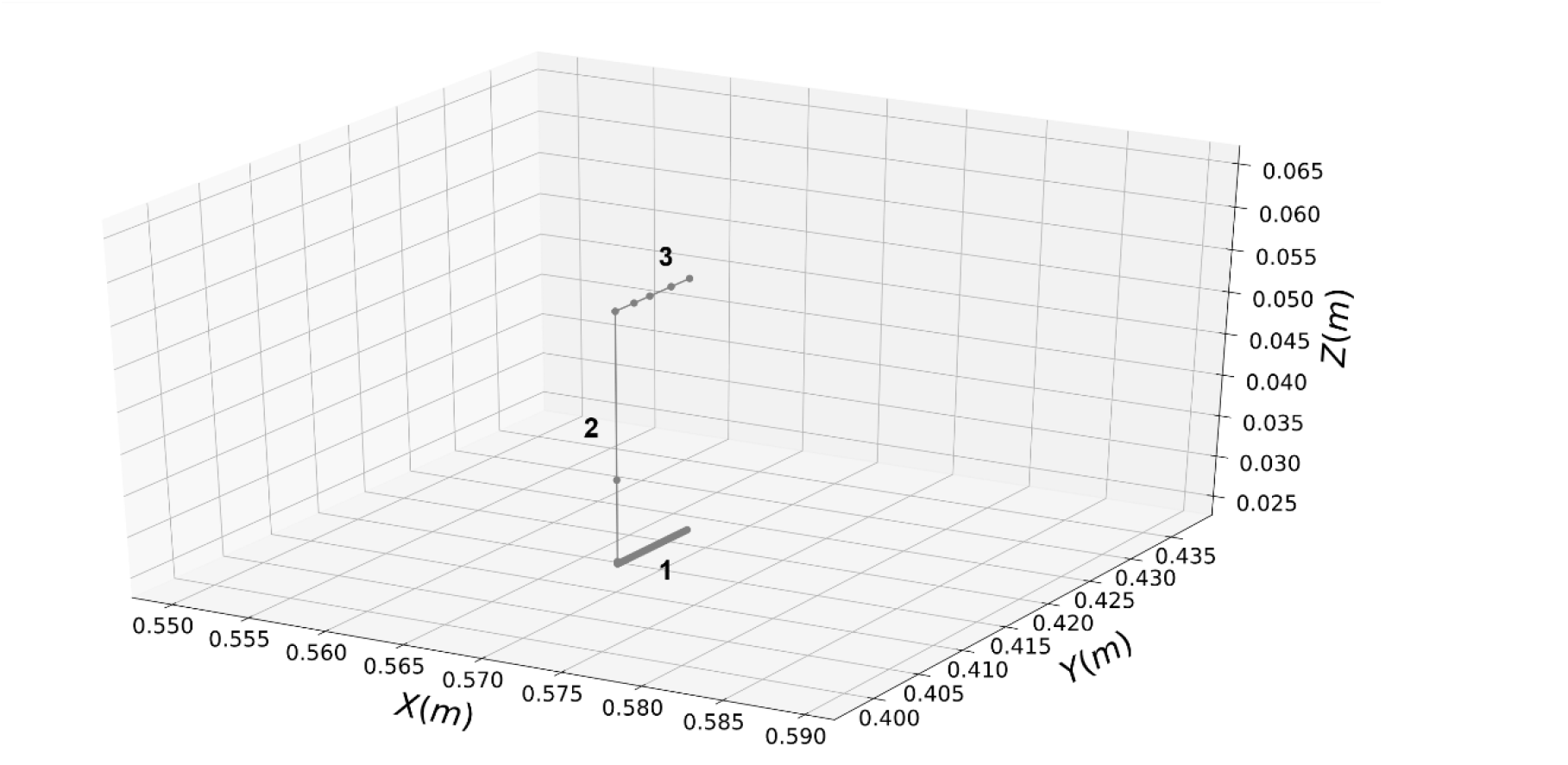
The trajectory generated for the optimization of robotic motion towards volumetric printing.

A volumetric AM robotic system should follow trajectories deriving from G-code files. However, volumetric approaches in small-scale manufacturing, from a trajectory generation point of view, are few [29-31]. Also, some of these approaches are either not open-source or very tailored for a specific hardware. One of the RAVEN platform goals is to provide a universal platform for robotic-based AM, meaning that the software platform can be easily implemented when the hardware changes.

To control the whole system and generate appropriate trajectories, more parameters need to be set and controlled compared to a conventional 3D Printing system, in which the path the extruder follows is planar and G-code-dependent. The most important parameters defined in a G-code file are the Cartesian coordinates of each point in the trajectory and the material deposition rate for that segment of motion.

An *ad hoc* slicing procedure is necessary in order to fully exploit the robot’s redundant DOF and all the potential that comes with it. The final goal of the new 7-DOF AM apparatus is the achievement of volumetric printing of biomedical grade polymers. Thus, complex architectures will be achieved easily, maintaining the continuity in fiber deposition and avoiding collisions of the printer-head with the construct manufactured in previous steps.

Commercial and free-to-use slicing software do not offer the possibility to customize the slicing process. Objects are sliced along the z-axis in a 2D and planar fashion (i.e., setting the thickness of each layer). Furthermore, volumetric slicing is pretty uncommon in small scale manufacturing. A first version of a non-planar and volumetric custom-made slicer software has been developed using the Grasshopper (GH) plugin for Rhinoceros (Robert McNeel & Associates, Seattle, WA, USA) and employed for both the design phase (e.g., to design complex and nature-inspired architectures) and the G-code generation, which was then further processed towards the robotic application.

### 2.4 Extruder integration

As previously mentioned, the system is composed of the robotic arm and (so far) an FDM filament extruder. The coupling between these two components occurs through custom-made supports, which were 3D printed as well. To improve the trajectory planning, the design of the extruder was incorporated into the robot configuration file, which is used for the planning process itself, by manually editing the original file, which is provided by the manufacturer company.

The choice of that specific extruder was crucial to allow for an easy control of all printing parameters in the early stages of the development. In particular, it is controlled through the Marlin firmware, which runs on the control board of the system. In general, every G-code is used in two different and simultaneous processes.

In order to synchronously send commands to the extruder while the robot is in motion, a method to send the G-code to the extruder via serial communication was developed. This method relies on the Descartes planners ability to move between points with different velocities according to the variation of the Time Between Points (TBP). This multithreading approach was used to run two programs synchronously, one to control the robot and one to control the extruder. In particular, when the robot started moving, another thread was used to run a program, which sent the G-codes to the extruder. Here, all the necessary calculations, which are part of the trajectory planning algorithms, are done in advance to prevent any delays that could interfere with the printing process and then the commands are sent synchronously to the robot and the extruder. Furthermore, to allow for a fast cooling of thin filaments in air, an advanced cooling system was implemented through typical 3D printing fans and a custom-made fan duct.

### 2.5 Assessment of extrusion methods

To test the capability of the RAVEN AM apparatus, printing tests were performed extruding both a commercial (Jupiter Serie, 123-3D ink, Netherlands) and pellets (NatureWorks, Resinex Netherlands BV, Netherlands) of PolyLactic Acid (PLA). In particular, the latter were processed through a twin extruder (HAAKE™ MiniCTW Micro-Conical Twin Screw Compounder, Thermo Fisher Scientific, Waltham, MA, USA) at 200°C and 80 rpm to turn them into filaments with a diameter of 1.75 mm.

The robotic RAVEN system was first tested as a conventional extrusion-based planar AM platform. A few test prints were performed with the design of a simple 0/90° scaffold for RM at different feedrates (FR), namely 2.4, 3.2, 4, 8, 12, 16, 24, and 32 mm/s, to check the influence of the system vibrations on the final resolution of the fabricated parts. In particular, the dimensions of the scaffolds were 10×10×10 mm, while Strand Distance (SD) was set to 800 µm, thus setting the desired value of Filament Gap (FG, i.e., porosity) to 400 µm.

Then, the ability of the Descartes planner to specify the end-effector configuration for each of the waypoints was put into use. Initial trials were done by printing on a tilted surface. A more precise control on the robot joints movements allowed for the fabrication of non-planar geometries, which paved the way to more complex volumetric samples, that required more control on the end-effector rotation angle. First, arch-like structures were printed to test the volumetric approach as a combination of planar and “in air” printing. A first vertical sine-like was designed and tested to assess the ability of the RAVEN system to rotate its end-effector around the three axes.

## 3. Results

### 3.1 Optimization of motion planning

The motion planning optimization results are showed in Figure 4. The robot had to follow the cylindric trajectory presented in Figure 2. The MoveIt Cartesian planning method did not result as the best for small motions, thus its applicability towards the RAVEN system application as well (Figure 4a). Furthermore, it lacks the capability to control the velocity and acceleration of the robot. To overcome this issue, an alternative solution based on the IK service provided by the Intera robotic platform (Rethink Robotics) was implemented to enable a control over joints velocity. However, this resulted in non-continuous movements between one segment and the other. In particular, the robot stopped for some seconds after each movement. This pause caused a large deviation from the desired trajectory, as shown in Figure 4b. Furthermore, many vibrations were recorded in the trajectory.

**Figure 4:**
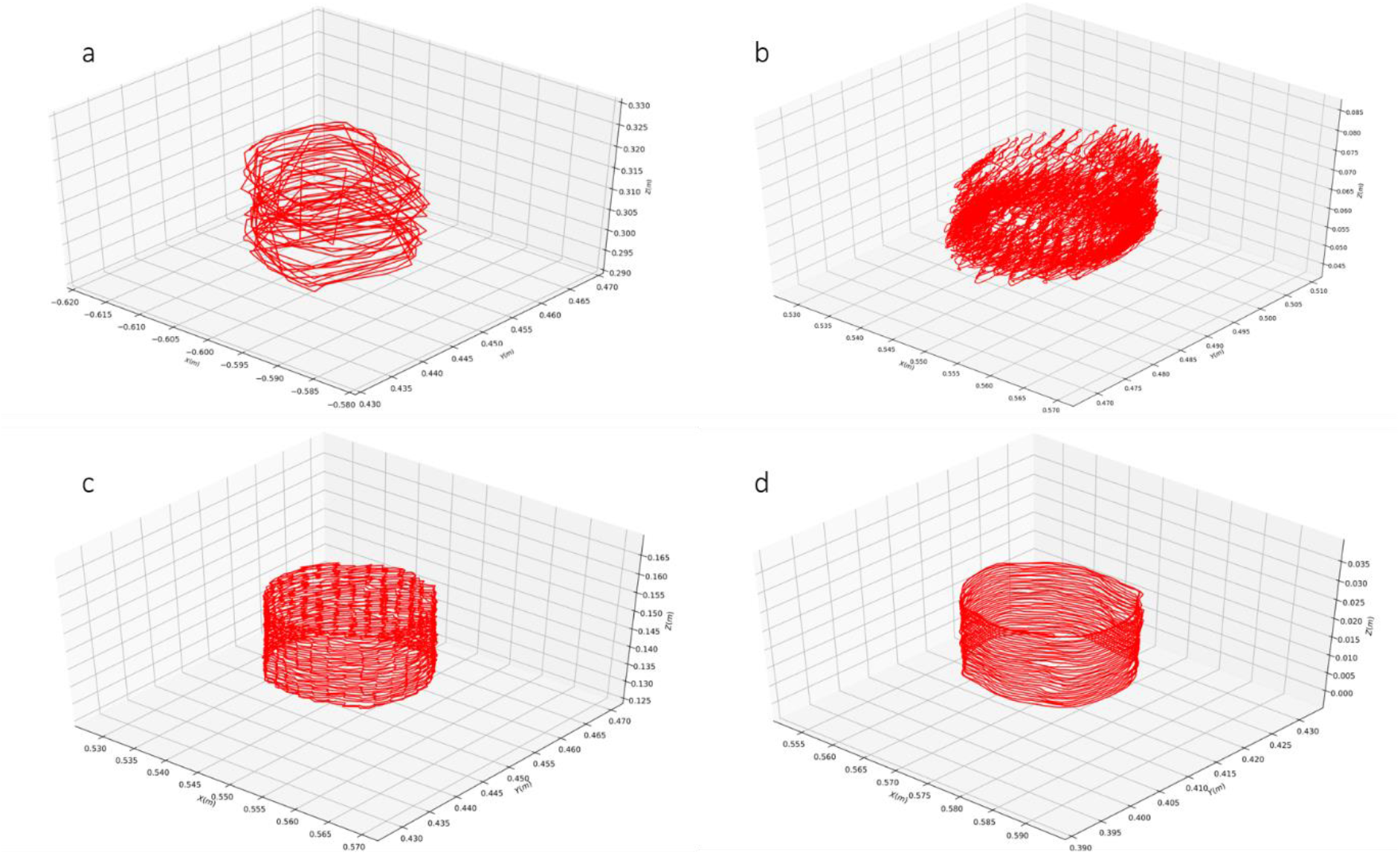
Optimization of motion planning for different planners/IK solvers in terms of trajectory fidelity. a) MoveIt Cartesian Planning + KDL; b) MoveIt Cartesian Planning + IK Intera solver; c) PILZ Planner + KDL; d) Descartes Cartesian Planner + IKFast IK Solver.

PILZ Industrial planner is mainly used for industrial robots and large-scale trajectories. However, it allows for Cartesian motion of the end-effector. PILZ-based algorithms allowed for a better individual control on each motion parameter for each trajectory segment, compared to MoveIt Cartesian Planner. The robot motion deriving from PILZ planning resulted smoother and more repeatable than the first option, both in the simulated and real environment. However, such approach does not allow for control over joints velocity for each motion segment, instead only on the whole trajectory. Furthermore, the small-scale application of this planner resulted in the formation of jerky motions, as shown in Figure 4c.

Descartes allows for the control of the end effector configuration in space along specific trajectories, which are defined through Cartesian points. As an added advantage, it is possible to control the speed of the end effector individually using TBP as a control parameter. This is a crucial requirement to develop a robust extrusion-based small-scale AM platform. This approach improved the quality of the planned trajectories, as shown in Figure 4d. Furthermore, the stability of the system improved in terms of vibrations, as shown in Figure 5.

**Figure 5:**
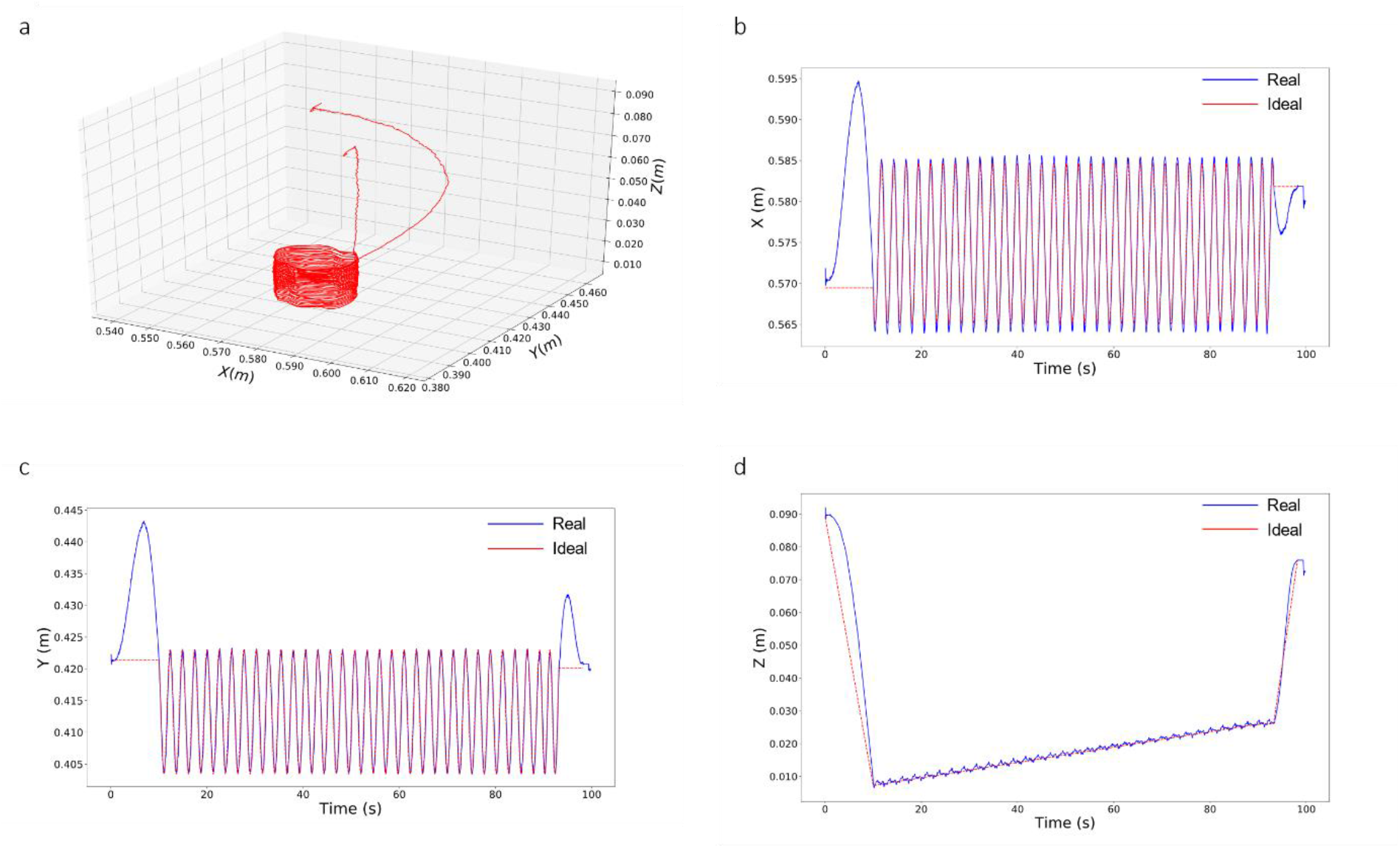
Trajectory fidelity of Descartes Planner towards small-scale movements in terms of planned vs. actual end-effector movements. a) the cylindrical trajectory the robot followed; b) extraction of end-effector movements along x-axis; c) extraction of end-effector movements along y-axis; d) extraction of end-effector movements along z-axis.

The Descartes planner showed high trajectory fidelity, also in terms of joints vibration (i.e., end-effector) in the three directions. The misalignment along the x and z direction was ± 1mm. Such values, which are due to hardware limitations and could seem significantly high for small-scale manufacturing purposes, can be compensated through further optimization of printing parameters, such as feedrate, flowrate and cooling.

### 3.2 Trajectory generation and planning

Results from the first trials of trajectory optimization towards volumetric AM are showed in Figure 6 and Figure S1. In order to test the 7 DOF capabilities of the robotic system, vertical trajectories were generated and tested at different feedrates. The robot trajectory deviated from the desired one for slow movements and if the points density was too small. Thus, a minimum requirement of 4 points/mm was set towards an accurate trajectory planning. Very fast feedrates resulted in jerky movements as well. Furthermore, the maximum printing feedrate of the extruder (i.e., 150 mm/s, which was provided by the supplier) was kept into consideration.

**Figure 6:**
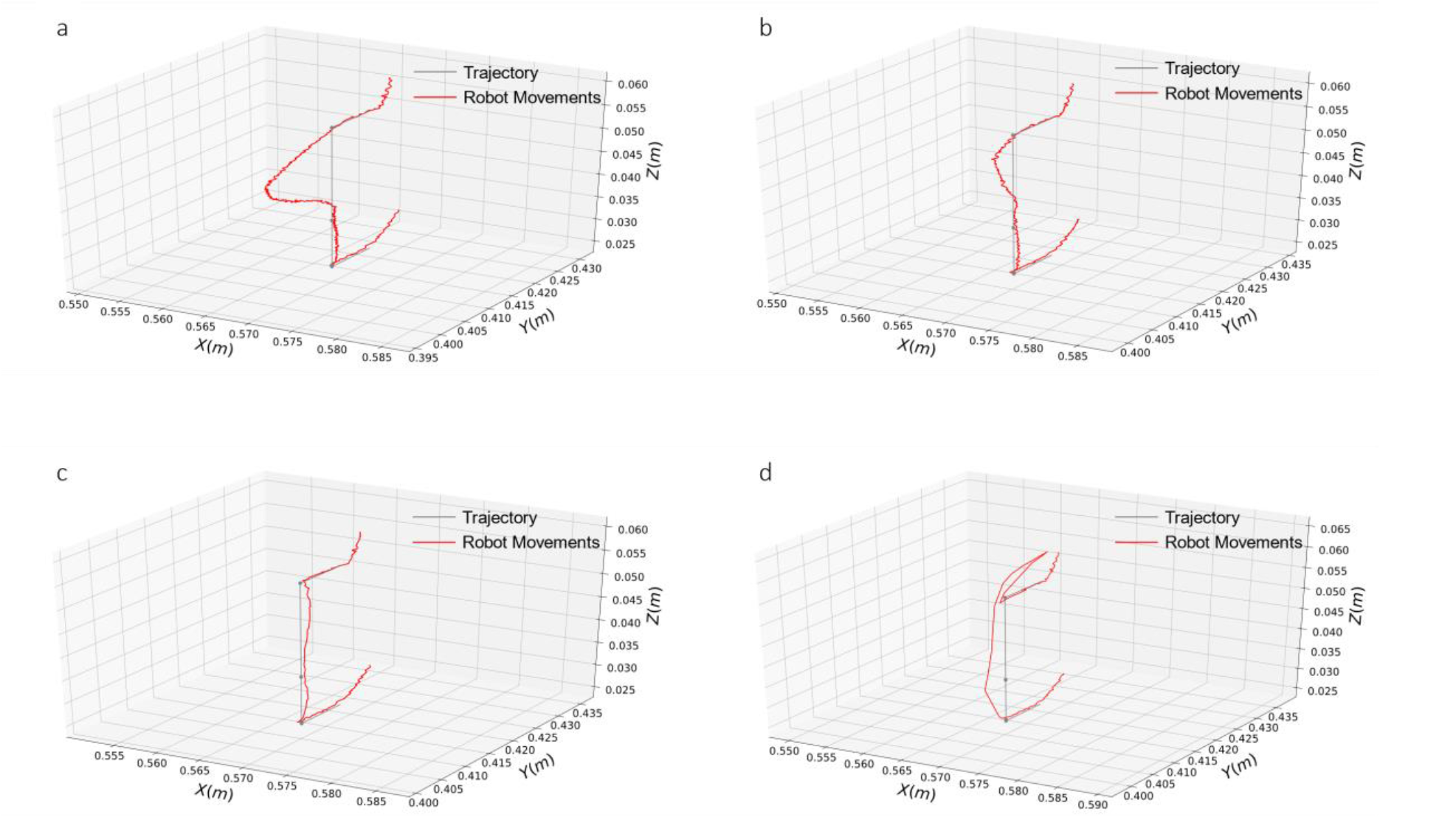
Results of trajectory optimization towards volumetric AM for different FR values. a) 1.6 mm/s; b) 4 mm/s; c) 24 mm/s; d) 160 mm/s

Since in a 7-DOF robotic system the end-effector orientation and rotations around the three axes can vary point-by-point, more parameters are required within the same code. This is why a new code which incorporates these new parameters was developed. In particular, the initial G-code was post-processed to include the new information (i.e., orientation of the end-effector with respect to the x,y, and z axis).

The requirement of extrusion of continuous lines and segments along which the robot has to move, combined with the cooling times and change of direction during printing, calls for additional parameters to be included. In particular, a new parameter, which was defined as “segment number”, was introduced in the new G-code based file. This parameter acts as a splitting flag between two consecutive segments in the trajectory generation phase. Nonetheless, the continuous extrusion was guaranteed.

A graphic representation of the RAVEN software algorithm is presented in Figure 7, where the simultaneous execution and data exchange between the trajectory planning and extrusion pathways is shown. In particular, the integration of TBP, which is extracted from the segments generation during the trajectory planning, is integrated into the “extrusion G-code” to recalculate feedrates and E values (i.e., the amount of material to be extruded between two consecutive points).

**Figure 7:**
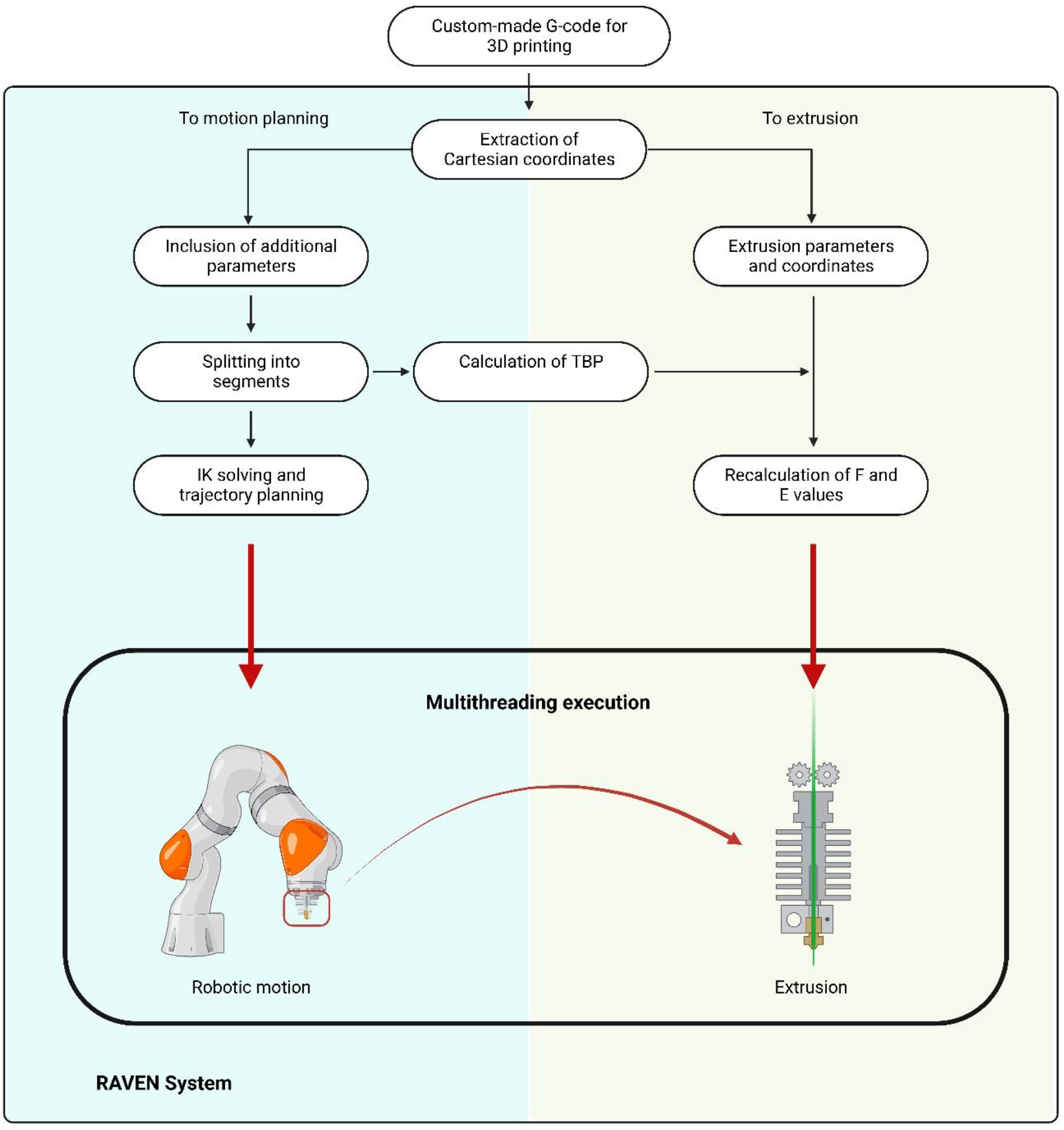
Representation of the RAVEN system software workflow

### 3.3 Extruder integration

Following the RepRap philosophy, the rationale of the hardware integration was to 3D print the custom-made pieces for the new AM apparatus, which has the capability to print its own replacing parts or upgrades in the future. All the designs are shown in Figure 8. In particular, the custom-made extruder stand (Figure 8b) was designed in order to have a precise alignment between the nozzle tip axis (on the extruder) and the z-axis of the end effector of the robot. This way, a more accurate control over robot movements was possible. The fan stand and fan ducts (Figure 8c and 8d) were designed in order to be compatible with the aforementioned extruder stand. To do so, everything was designed around the extruder design in SolidWorks (Figure 8a).

**Figure 8:**
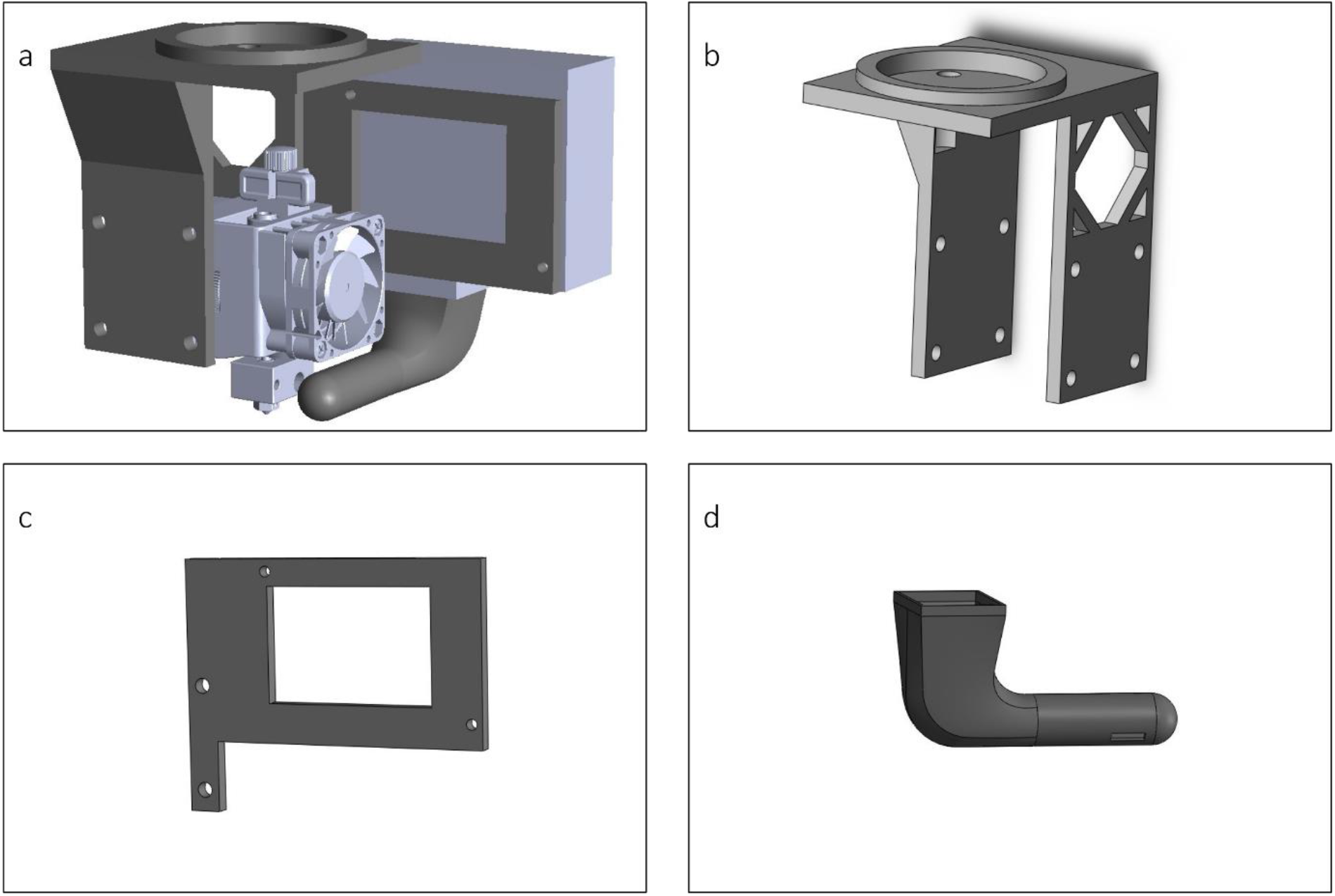
3D models of the custom-made supports for the robot-extruder interface. a) The final assembly design; b) Extruder stand: c) Fan stand; d) Fan duct.

The custom-made code to manipulate the G-code and send synchronous signals to the robot and the extruder was successfully tested and implemented.

### 3.4 Assessment of extrusion methods

Printing results of the RAVEN system are showed in Figures 9-11. As previously said, the system was initially tested in a planar way, and thus typical 3D porous scaffolds for RM were designed and sliced in the custom-made platform and then extruded (Figure 9a) for different FR values. The system followed the trajectory in a precise and repeatable way.

**Figure 9:**
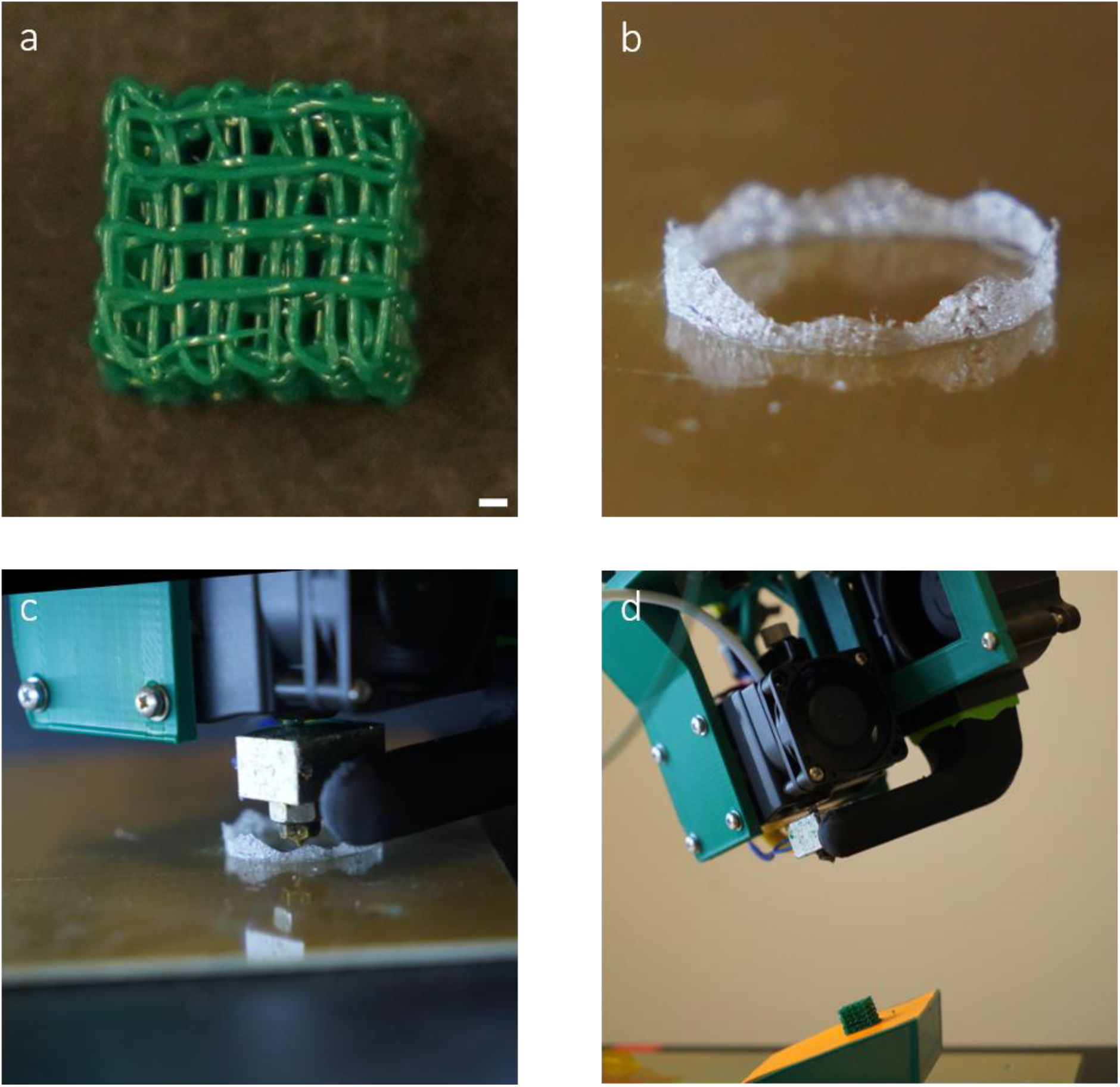
Planar and non-planar extrusion capabilities of the RAVEN AM system. a) Printed planar scaffolds for RM. Scale bar is 1 mm; b-c) Results of the non-planar extrusion assessment; d) Printed devices on a tilted surface.

The final resolution of the samples was lower in the case of high and slow FR. In particular, for feedrates of 2.4, 3.2, and 4 mm/s, a difference between the first layer and the other layers was evident in terms of Filament Diameter (FD) (i.e., 200 µm), whilst for FR of 8, 12, and 16 mm/s there was no major difference between layers (∼ 60 µm). Vibrations, which are due to robotic hardware limitations (i.e., joints springs), were not a major issue for FR values of 8, 12, and 16 mm/s, as the FD fluctuated in a range of 70, 40, and 80 µm from the nominal (i.e., 400 µm), respectively. In terms of pores dimension and morphology, the slight misalignment of the robot in the x-axis resulted in the formation of non-homogeneous pores, thus the Filament Gap (FG) was not constant, but fluctuated between smaller values (around 400 µm with a deviation of 75 µm for all the FR combinations) and higher values (up to 1600 ± 120 µm).

When the planar capabilities of the robot were assessed, non-planar motions were tested. In particular, a gradually non-planar geometry was designed within the custom-made designing/slicing platform. Successful trials (Figure 9b-c and Video S1) proved the capability of both the slicing platform and the RAVEN system to switch from a 2.5D to a proper 3D *xyz-based* printing, in the way the points for the G-code are generated and the extrusion occurs, respectively. This capability is crucial when aiming to *in situ* applications, in the presence of non-planar surfaces and in order to follow non-planar trajectories, such as the precise alignment of collagen fibers in specific tissues. In order to test the RAVEN potential to work in a volumetric way while changing its end-effector (and thus, extruder) orientation during extrusion of continuous lines, some initial extrusion trials on tilted surfaces were performed (Figure 9d and Video S2). In particular, exploiting the potential of the custom-made slicer, a planar design was projected onto a surface with a specific angle. The tilted platform was 3D printed as well and the robotic system was programmed to have its end-effector oriented by the specific aforementioned angle.

Volumetric printing was first approached as a combination of planar and “in air” printing, through the arch-like structures showed in Figure 10 and Video S3, which were printed both as a single layer (Figure 10a-b) and as multiple, intertwining layers (Figure 10c-d). The advanced and custom-made cooling system allowed for a fast cooling of the extruded filaments which did not collapse. Such structures pave the way for advanced *in situ* scaffolds fabrication.

**Figure 10:**
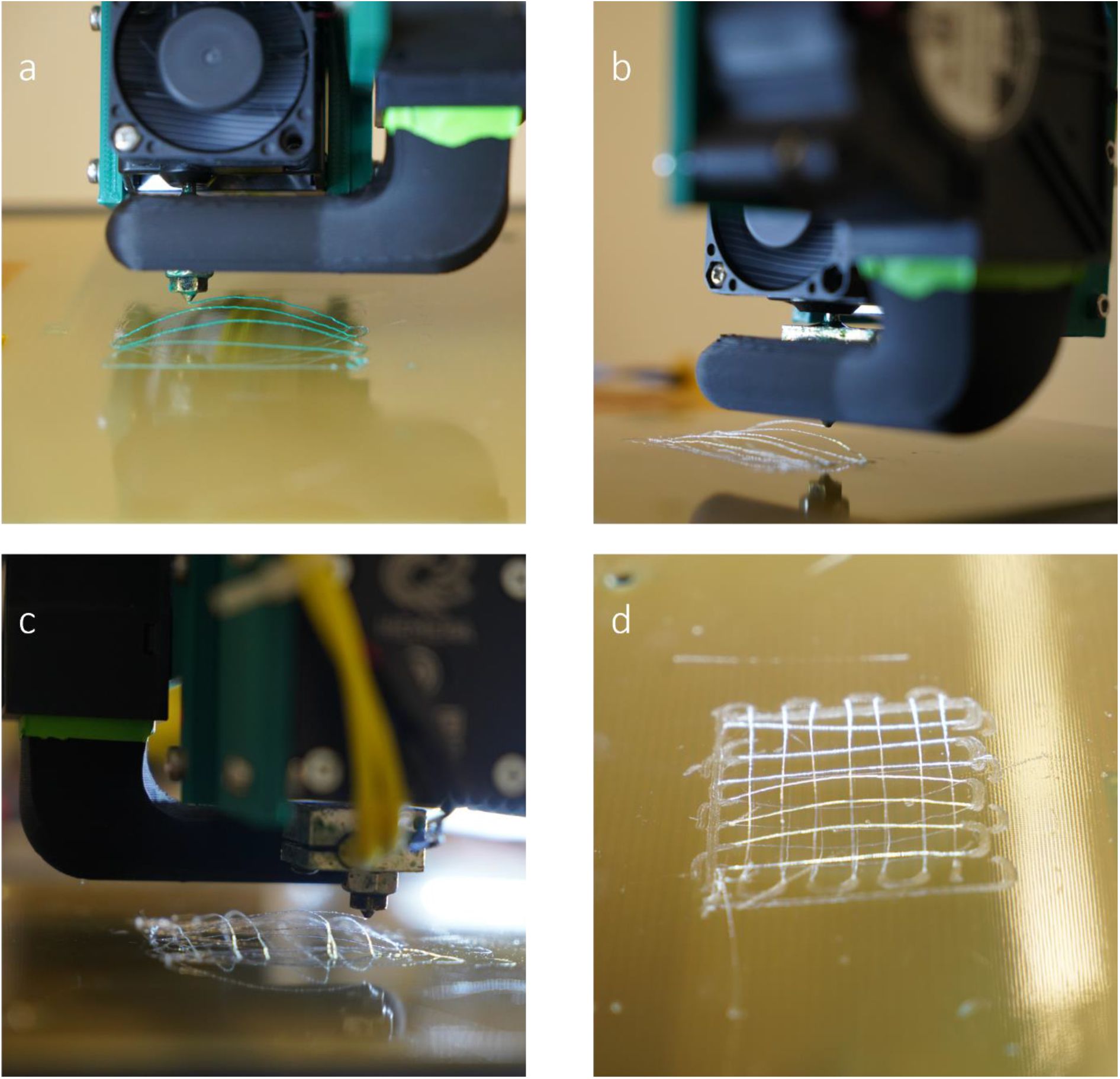
Arch-like structures. a-b) Extrusion of single-layer arch-like structures; c-d): Extrusion of multi-intertwining-layers arch-like structures.

Proper volumetric and in air printing was tested combined with the capability of the robot to change direction during motion. The successful extrusion of the vertical sine-like designs proved that, as shown in Figure 11 and Video S4. Such complex extrusion patterns once again corroborate the ability of the RAVEN system to follow precise trajectories, taking full advantage of its redundant DOF (i.e., the 7^th^ one). Furthermore, the FD of the extruded sine waves was 0.408±0.014 mm, which, compared to the 0.4 mm diameter of the needle tip (and thus, nominal FD), proved the efficiency of the cooling system and how the interplay of different parameters, such as feedrate and printing speed, can overcome the effect of hardware vibrations in the final device.

**Figure 11:**
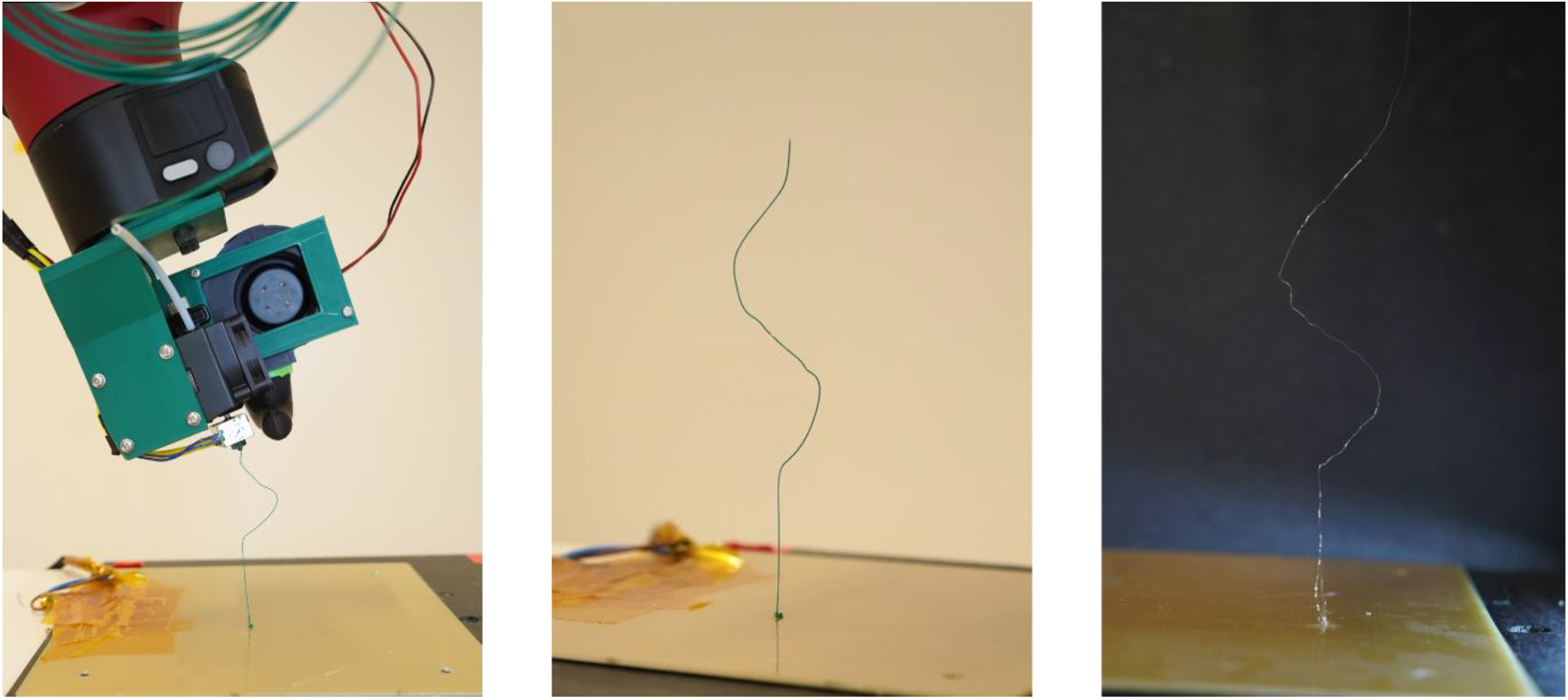
Extrusion of sine-wave designs.

## 4. Discussion

As the need for devices that better and better mimic native tissues increases, a switch to a volumetric fabrication methodology of such devices is required as well. The RAVEN system aims to fill such gap in the RM field, providing a universal platform in terms of software and hardware integration, which is transferable and custom-made. Since RAVEN finds its place in a very specific field, namely small scale Additive Manufacturing, robot motion and trajectory generation had to be optimized towards this goal.

Starting from known results in the robotic community for large scale motions and applications, different tests were conducted to find the most suitable combination of Cartesian planner and IK Solver for the RAVEN system. The Descartes planner, together with the IKFast IK Solver, was selected in this context. Its capability to precisely follow trajectories deriving from Cartesian points, as well as to control motion parameters for every sub-segment of the trajectory, makes it one of the best options for 7 DOF-based AM, in which precise fibers directions needs to be achieved. Furthermore, towards *in situ* applications where obstacle avoidance and environmental knowledge are crucial, full control on all the joints is needed. IKFast allows for the exploitation of all the DOF and more complex motions. Trajectory fidelity analyses resulted in a slight misalignment along the x and z axes, which could be compensated in the future with further tests. *Fortunato et al*. [32] achieved this with a 5DOF robotic system, for non-planar *in situ* bioprinting of gels, for skin regeneration applications. To tackle such applications for volumetric extrusion-based AM, further compensations of movements should be assessed, as trajectory generation and motion planning tests towards small-scale AM showed. Furthermore, thermoplastic medical-grade polymers and extrusion-based applications increase the complexity of the whole system.

Applications in literature involve less universal implementations, based on more commercial robots which can be easily used through simulation/actuation software (i.e., MATLAB) [32, 33]. The development of a custom-made control algorithm in ROS-Linux environment will make the RAVEN system hardware/software-free, as it will be easy to adapt it to control any 5/6/7-DOF robotic arm and FDM serial-based extruder. Furthermore, the RepRap rationale makes the hardware interface open source and accessible.

The RAVEN system was tested as a proper AM apparatus step by step, from planar to full volumetric approaches, going through more and more non-planar designs. A good trade-off between points density and FR was achieved, especially for vertical trajectory segments. However, when planar tests were performed, lack of precision and a slight misalignment in the x direction was recorded and quantified. When aiming to scaffolds for RM, deposition fidelity, as well as higher fidelity in the FD are crucial. *Bin Ishak et al*. [18] developed a system based on an industrial 6-DOF robot, which was controlled with a closed system and an FDM extruder to print planar structures through multi-plane toolpaths. Similar approaches could be useful to address the fidelity issue of the RAVEN system.

The non-planar capabilities of the system were successfully tested. Similarly to other approaches in literature [33], planar objects were projected onto non-planar surfaces and printed. In the case of the RAVEN AM technology, these tests were useful to assess the control on the joints angles and end-effector orientation towards more complex applications. Furthermore, non-planar structures “in air” (i.e., the arch-like designs) were successfully printed and proved an interesting starting point for sectorial slicing devices, in which structural requirements (e.g., specific alignment of fibers) could change from one sector of the design to the other. In particular, the specific design of those multilayer structures, in which every other two layers are slightly shifted on the x direction, allowed for further test on the system capability to be reproducible and precise even for close starting points.

A recent work from *De Marzi et al*. [34] was focused on the development of a robotic platform for UV-DIW printing, made of a 6-DOF robot and an extruder. There, vertical segments were printed in a discontinuous way, to allow for cross-linking to occur. RAVEN allows volumetric printing, taking full advantage of the associated custom-made cooling system, which enabled the extrusion of continuous lines without under or over extrusion. The change of direction in the vertical sine-wave design allowed for a more complex and gradual control on the end-effector angle during the movements, which is a critical requirement for obstacle avoidance algorithms and applications.

## 5. Conclusions

In the context of extrusion-based AM towards advanced RM, the RAVEN system was developed and tested as a new AM apparatus. The software and hardware interface between the 7-DOF robotic arm and the FDM extruder is custom-made, following the RepRap concepts, and transferable to other hardware.

Many tests were performed to select the best Cartesian planner as well as IK solver, in terms of trajectory fidelity. Modifications to traditional G-codes for AM were necessary to allow for a synchronous execution of robot motion and extrusion, which were possible through a multithreading approach.

Planar, non-planar, and volumetric extrusion tests were performed to assess the system capabilities in terms of AM trajectories fidelity and full exploitation of the redundant DOF. Such tests showed the potential of the system towards volumetric printing of complex structures fabrication for RM, where precise control over fibers alignment and direction is needed.

Limitations of this system can be identified in the jerk of specific movements, which are due to intrinsic limitations of the hardware when performing small movements, but that can be overcome through further optimization of printing and motion parameters. Furthermore, an FDM extruder could not be the best choice when switching to specific medical grade biopolymers which are typically extruded through pneumatic-based extrusion.

Future work will include more optimization of the volumetric printing, in order to achieve more and more complex structures easily, as well as a more robust platform for the software interface between the robot and the extruder. Furthermore, other extrusion tools (e.g., pneumatic based extruder heads) will be included in the RAVEN system, thus their integration in the hardware/software interface in terms of coupling and control.

## Acknowledgements

This publication is part of the project 3D-MENTOR (with project number 18647) of the VICI research programme, which is financed by the Dutch Research Council (NWO).

