## Supplementary material for "RAVEN: development of a novel volumetric extrusion-based system for small-scale Additive Manufacturing": Figure S1

**Supplementary Figures**

*Figure S1: Results of trajectory optimization towards volumetric AM for different FR values. a) 0.4 mm/s; b) 0.8 mm/s; c) 0.32 mm/s; d) 1.6 mm/s; e) 2.4 mm/s; f) 3.2 mm/s; g) 4 mm/s; h) 8 mm/s; i) 16 mm/s; j) 24 mm/s; k) 32 mm/s; l) 40 mm/s; m) 80 mm/s; n) 160 mm/s.*

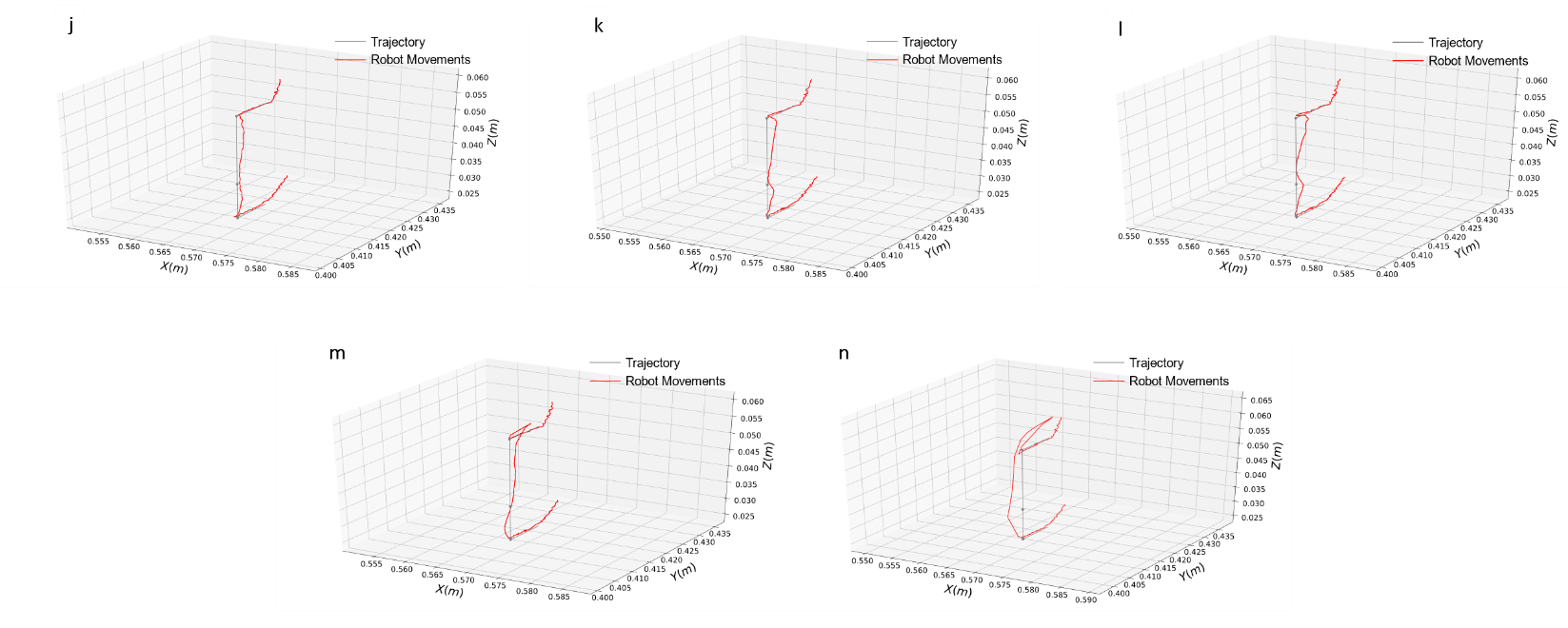

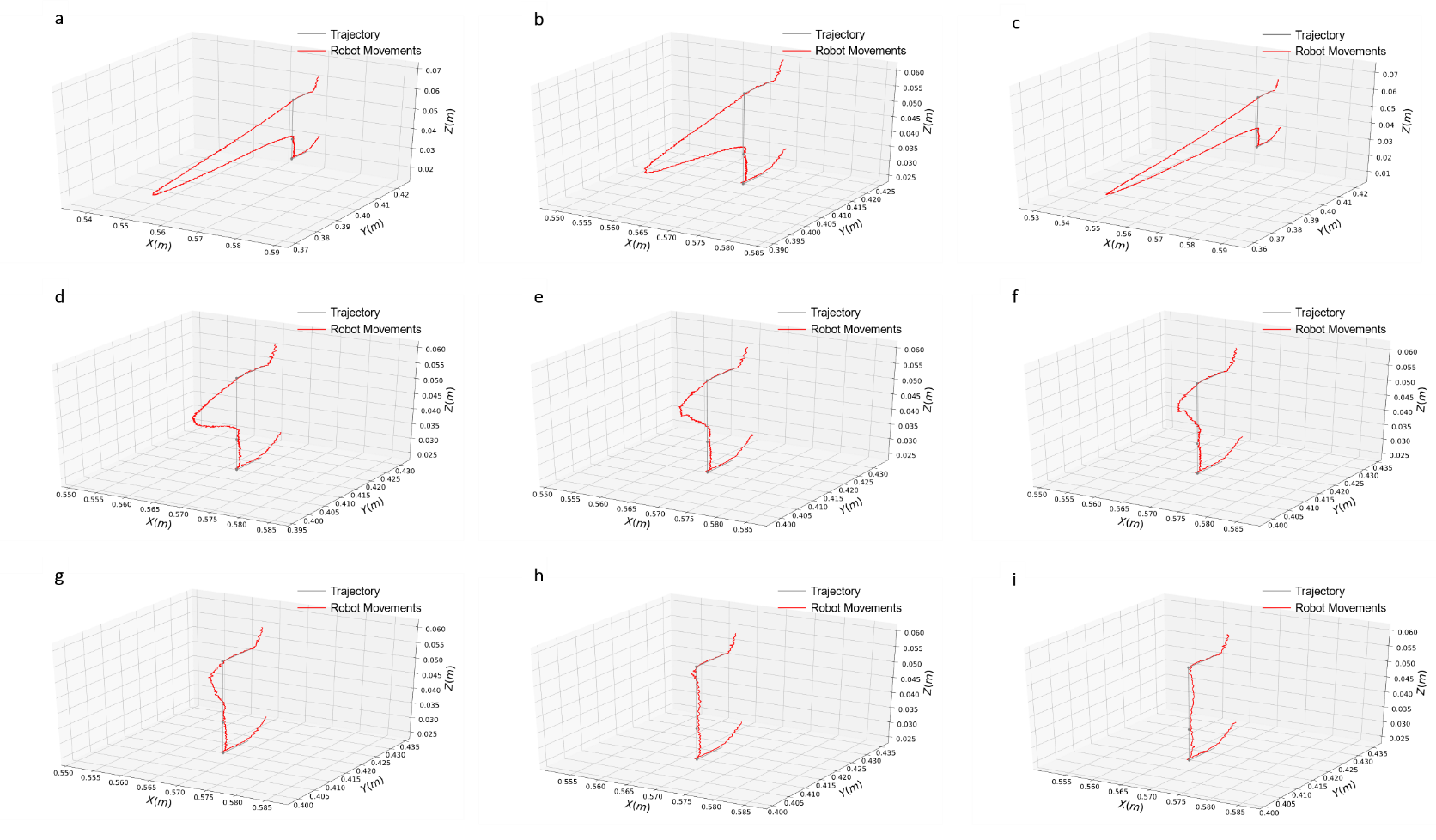
